# Preponderance of Enterovirus Species C in RD-L20B cell culture negative stool samples from children diagnosed with Acute Flaccid Paralysis in Nigeria

**DOI:** 10.1101/116004

**Authors:** J.A. Adeniji, A.O. Oragwa, U.E. George, U.I. Ibok, T.O.C. Faleye, M.O. Adewumi

**Author notes:** AUTHOR NAME and EMAIL. ADENIJI, Johnson Adekunle, 0000-0002-0406-6707(ORCID). ORAGWA, Arthur Obinna, GEORGE, Uwem Etop, IBOK, Ukeme Ibanga, FALEYE, Temitope Oluwasegun Cephas, 0000-0001-7706-9493 (ORCID). ADEWUMI, Olubusuyi Moses, 0000-00025172-5808 (ORCID).

## Abstract

Recently, a reverse transcriptase seminested polymerase chain reaction (RT-snPCR) assay was recommended by the WHO for direct detection of enteroviruses from clinical specimen. In this study, we use the assay and its modification to screen acute flaccid paralysis (AFP) samples previously confirmed negative for enteroviruses by the RD-L20B algorithm.

Thirty paired stool suspensions collected in 2015 as part of the national AFP surveillance program in different states of Nigeria were analyzed in this study. The samples were previously confirmed negative for enteroviruses by the polio laboratory in accordance with the WHO recommended RD-L20B cell culture based algorithm. Two samples previously confirmed to contain enteroviruses were included as positive controls. All samples were subjected to RNA extraction, and the RT-snPCR assay and its modifications. All amplicons were sequenced and enteroviruses identified using the enterovirus genotyping tool.

Overall, amplicons were recovered from the two controls and 50% (15/30) of samples screened. Fourteen were successfully typed of which, 7.1% (1/14), 21.4% (3/14), 64.3% (9/14) and 7.1% (1/14) were EV-A, EV-B, EV-C and a mixture of EV-B and C (EV-C99 and E25), respectively. The two controls were identified as EV-C99 and CV-A1, both EV-Cs. The PV-2 detected had VP1 ILE143. Hence, a vaccine strain.

The results of this study showed that about 50% of enterovirus infections (including some Sabin PV2s) are being missed by the RD-L20B cell culture based algorithm. This highlights the value of the RT-snPCR assay and its modifications. The circulation and preponderance of EV-Cs in Nigeria was also confirmed.

## Introduction

Enteroviruses are members of the genus *Enterovirus* in the family *Picornaviridae,* order *Picornavirales.* The type member of the genus is poliovirus, the etiologic agent of poliomyelitis and a member of Enterovirus species C (EV-C); one of the twelve species in the genus [1]. EV-B has most (>60) of the enterovirus serotypes that have been described till date [1]. Classically, serotypes were defined using neutralization assay [2, 3]. More recently, a correlation has been found between the nucleotide sequence of the VP1 gene and serotypes [2, 3]. Consequently, the nucleotide sequence of the VP1 gene is being used for enterovirus serotype/genotype designation, and has allowed the discovery of several new types [1].

Majority of the nonpolio enteroviruses (NPEVs) isolated till date were isolated courtesy the poliovirus eradication effort (Global Polio Eradication Initiative [GPEI]). The GPEI was inaugurated in 1988 via a World Health Assembly (WHA) resolution to eradicate poliovirus [4]. The effort has centered on intensive vaccination (with both oral and inactivated polio vaccines [OPV and IPV]), and surveillance (for poliovirus in children <15 years old with acute flaccid paralysis (AFP) and sewage contaminated water) [5-7].

As part of the surveillance effort is a team of about 140 laboratories (Global Polio Laboratory Network [GPLN]) that are WHO certified and responsible for poliovirus laboratory diagnosis globally in accordance with the WHO guidelines [5, 6]. These guidelines have isolation of poliovirus in cell lines (RD and L20B) as the core on which the rest of the algorithm is built.

Though very sensitive for poliovirus detection and identification, studies have shown that the RD-L20B algorithm also selectively supports EV-B detection while not significantly supporting nonpolio EV-C (NPEV-C) isolation [8-10]. This suggests that some NPEVs are missed by the RD-L20B algorithm. Furthermore, we have previously shown [9] that by including a cell line like MCF 7 in the isolation algorithm, enteroviruses that had been missed by the RD-L20B algorithm can be subsequently recovered from sewage contaminated water (environmental) samples.

Recently [11], a reverse transcriptase semi-nested polymerase chain reaction (RT-snPCR) assay (first described by Nix et al., [12]) was recommended by the WHO for direct detection of enteroviruses from clinical specimen. The sensitivity of this assay has been confirmed by several studies [13-16] and we have further improved the assay by enhancing its enterovirus co-infection resolution capacity [16].

In this study, we use the WHO recommended RT-snPCR assay [11] and its modification [16] to screen AFP samples that had been previously screened, and declared negative for enteroviruses using the RD-L20B algorithm [6]. We show that about 50% of the samples had enteroviruses in them and EV-Cs were the predominant species recovered.

## Methodology

### Sample Collection and Processing

The samples analyzed in this study were obtained from the WHO National polio laboratory at the Department of Virology, College of Medicine, University of Ibadan, Nigeria. These stool suspensions were collected in 2015 as part of the national Acute Flaccid Paralysis (AFP) surveillance program in different states of Nigeria. The samples were previously analyzed in the polio laboratory in accordance with the WHO cell culture based algorithm recommended for poliovirus isolation [6] in order to determine the presence or otherwise of polioviruses and NPEVs in them.

Thirty cases whose samples showed no cytopathology in RD and L20B cell lines [6] and were consequently declared negative for enteroviruses were selected at random and analyzed in this study. The 30 cases consisted of 60 samples (two samples for each case collected within 24 hours). Each of the 30 paired samples was pooled (i.e. making 30 samples) and analyzed. In addition, two samples that had been earlier confirmed to contain enteroviruses were included to serve as positive controls.

### RNA Extraction and cDNA Synthesis

RNA was extracted from the thirty-two (32) samples using Jena Bioscience RNA extraction kit (Jena Bioscience, Jena, Germany) according to manufacturer’s recommendations. Similarly, cDNA was synthesized from the RNA extract using Script cDNA Synthesis Kit (Jena Bioscience, Jena, Germany) according to manufacturer’s recommendations. Firstly, 4.75μL of cDNA synthesis mix was prepared per reaction and contained 2μL of Script RT buffer, 0.5 μL of dNTP mix, 0.5 DTT stock solution, 0.5μL of RNase inhibitor, 0.25μL of SCRIPT RT and 0.25μL of each of the oligonucleotide primers AN32, AN33, AN34, and AN35 [11, 12]. Then, 5.25 μL of RNA extract was added to make 10 μL. This mixture was incubated at 42°C for 10 minutes followed by 50°C for 60 minutes in a Veriti thermocycler (Applied Biosystems, California, USA).

### Semi-nested Polymerase chain reaction (snPCR) Screens

One first round and three (PE, EV-A/C, EV-B) different second round PCR assays were done. For the first round PCR assay, the reactions were done in 30μL volumes. Each contained 6μL of Red Load Taq, 13.4μL of RNase free water and 0.3μL each of the forward (224) and reverse (222) primers, and 10μL of cDNA. A Veriti thermal cycler (Applied Biosystems, California, USA) was used for thermal cycling with the following conditions; 94^0^C for 3 minutes, followed by 45 cycles of 94^0^C for 30 seconds, 42^0^C for 30 seconds, and 60^0^C for 60 seconds, with ramp of 40% from 42^0^C to 60^0^C. This was then followed by 72^0^C for 7 minutes, and held at 4^0^C until the reaction was terminated.

All three second-round PCR assays were carried out with the first-round PCR product as template, with similar thermal cycling conditions except for the extension time that was reduced to 30 seconds. The same reverse primer (AN88) was used for all three second round PCR assays. The forward primers were AN89, 189 and 187 for Panenterovirus (PE), Enterovirus Species A or C (EV-A/C) and Enterovirus Species B (EV-B) PCR assays, respectively. Subsequently, all PCR products were resolved on 2% agarose gels stained with ethidium bromide and viewed using a UV transilluminator.

## Amplicon Sequencing and Enterovirus Identification

Amplicons of all the second round PCR assays with the expected band size (i.e. ~350 bp) were shipped to Macrogen Inc, Seoul, South Korea, for purification and nucleotide sequencing. Afterwards, the enterovirus genotyping tool (EGT) [17] was used for enterovirus species and genotype determination.

## Multiple Sequence Alignment (MSA)

Subsequent to enterovirus identification via the EGT [17], for nucleotide sequences generated from selected amplicons, multiple sequence alignments were done using reference sequences downloaded from Genbank and aligned using the CLUSTAL W program in MEGA 5 software with default settings [18].

## Nucleotide Sequence Accession Numbers

All the sequences generated in this study have been deposited in GenBank under accession numbers KY748282-KY748295.

## RESULTS

### PCR Result

The PE assay successfully amplified the expected ~350bp from 15 of the 30 (50%) samples screened in this study and the two controls. The EV-A/C assay successfully amplified the expected ~350bp from 11 of the 30 (36.7%) samples screened as well as the two controls. The EV-B assay successfully amplified the expected ~350bp from nine (9) of the 30 (30%) samples screened as well as the two controls. Altogether, amplicons were recovered from 15 of the 30 (50%) samples screened as well as the two controls (Table 1).

**Table 1:**
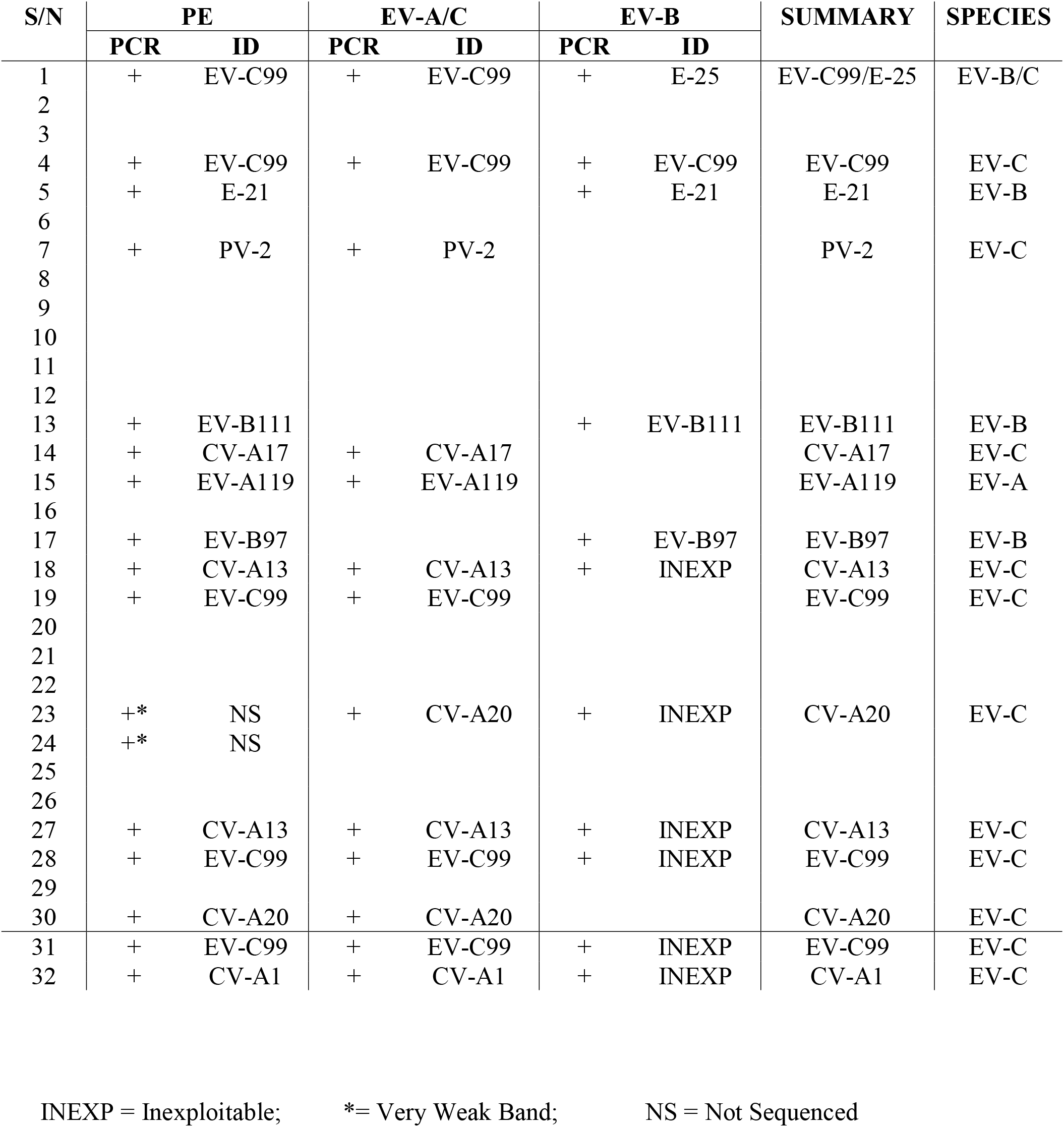
Enterovirus types detected and identified in this study

### Sequencing and Genotyping Result

For the PE assay, of the 15 amplicons generated, 13 were successfully sequenced alongside the two controls, while two (2) were not due to very weak bands. The isolates were identified as Enterovirus (EV) A119 (1 isolate), Echovirus-21 (E21) (1 isolate), EV-B97 (1 isolate), EV-B111 (1 isolate), Poliovirus 2 (PV-2) (1 isolate), CV-A13 (2 isolates), CV-A17 (1 isolate), CV-A20 (1 isolate) and EV-C99 (4 isolates), Specifically, of the 13 isolates detected by the PE assay, one, three and nine belong to EV-A, EV-B and EV-C respectively. The two controls were identified as EV-C99 and CV-A1, both EV-Cs (Table 1).

For the EV-A/C assay, all the 11 amplicons were successfully sequenced alongside the two controls, and identified as EV-A119 (1 isolate), PV-2 (1 isolate), CV-A13 (2 isolates), CV-A17 (1 isolate), CV-A20 (2 isolate), and EV-C99 (4 isolates). Of the eleven isolates detected by this study, ten belong to EV-C while one was EV-A. The two controls were identified as EV-C99 and CV-A1, both EV-Cs (Table 1).

For the EV-B assay, all the nine (9) amplicons and the two controls were successfully sequenced. However, sequence data from five of the isolates and the two controls were unexploitable due to the presence of multiple peaks. The remaining four isolates were successfully typed and identified as Echovirus 21 (E21) (1 isolate), E25 (1 isolate), EV-B97 (1 isolates) and EV-B111 (1 isolate). All the four identified isolates were EV-Bs.

### Co-infection resolution

Overall, these assays showed that only sample number one contained a mixture (EV-C99 and E25) and this was resolved (Table 1). It is however important to note that, had we relied on only the PE assay as recommended by the WHO, the presence of E25 in the sample would have been missed. Similarly, we might have missed CV-A20 in sample 23 (Table 1) had we relied on PE assay alone.

### Sequence based determination of the type of Poliovirus serotype 2 detected

Multiple sequence alignment (Figure 1) showed that the Poliovirus serotype 2 (PV-2) detected had the amino acid Isoleucine at the 143^rd^ position of the VP1 protein suggesting it to be a vaccine strain.

**Figure 1:**
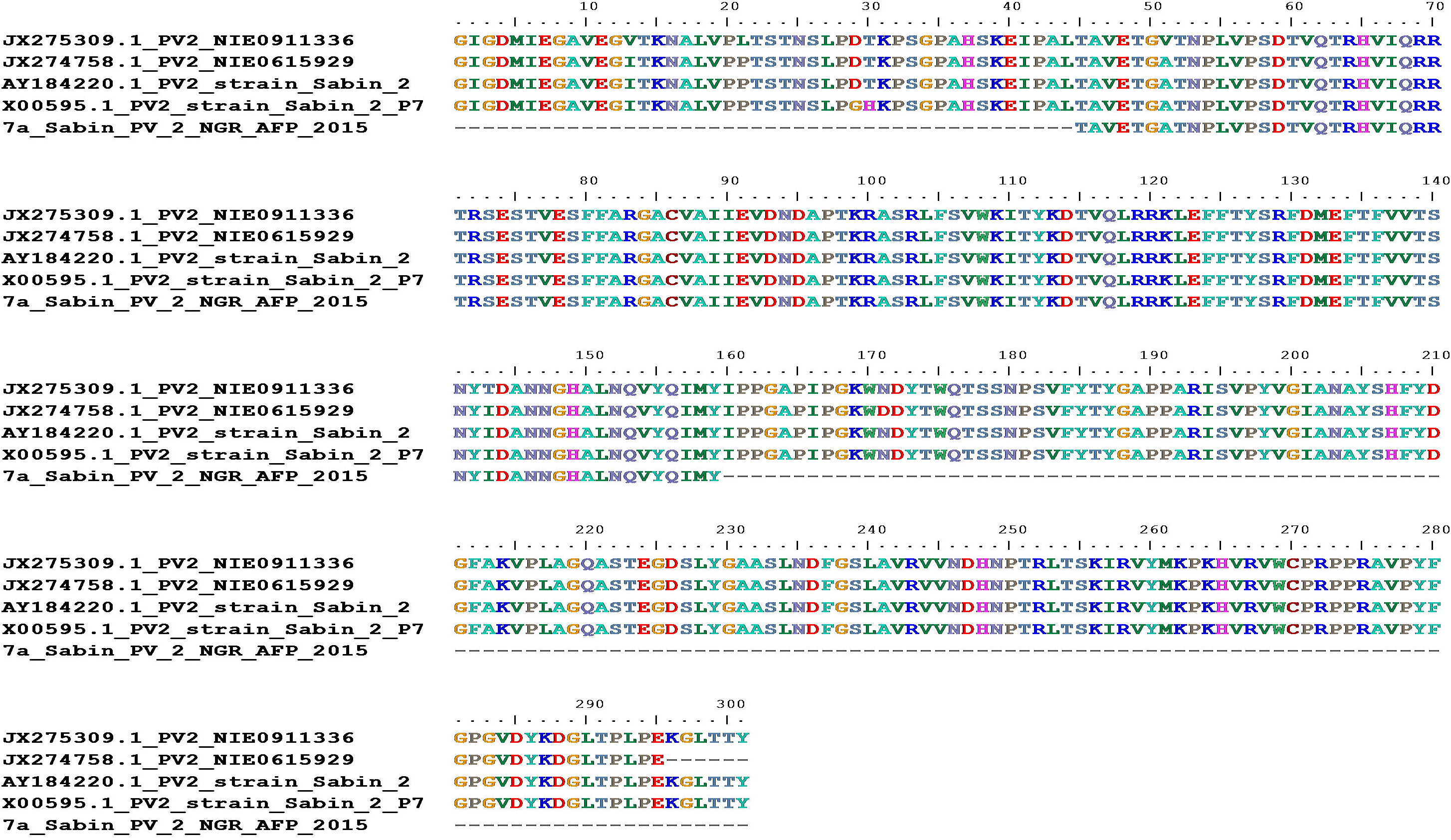
Alignment of the PV2 detected in this study (7a_Sabin_PV_2_NGR_AFP_2015) with some reference strains from GenBank. Note the Isoleucine in position 143 (I_143_) which indicates the isolate is a vaccine strain.

## DISCUSSION

In this study enteroviruses were detected in 50% (15/30) of the samples screened. However, enteroviruses could only be accounted for unambiguously in 46.7% (14/30) of the samples screened. Hence, the findings of this study suggest that about 46-50% of the stool samples from children <15 years old diagnosed with AFP in Nigeria that are declared negative for enteroviruses by the RD-L20B cell culture based algorithm might actually contain such. This suggests that a sizeable number of enterovirus infections is being missed by the RD-L20B cell culture based algorithm annually. The failure of this algorithm to detect some EVs may be due to factors like the cell lines not being susceptible and/or permissive to the EVs in question, presence of slow growing EVs in stool samples, poor handling of stool samples during transport to the laboratory and/or very low titre of viruses in the sample. We however understand that the detection limit of any PCR assay and the virus genome concentration in any sample both ultimately influence the sensitivity of the PCR assay. Hence, it is crucial to state that we are aware that the 46-50% detected in this study, might represent an underestimate.

Analysis of the enterovirus types found in this study revealed that 7.1% (1/14), 21.4% (3/14), 64.3% (9/14) and 7.1% (1/14) of the isolates were EV-A, EV-B, EV-C and a mixture of EV-B and C, respectively (Table 1). This study therefore confirms the circulation and preponderance of EV-Cs in Nigeria. Other studies where cell lines with EV-C ‘bias’ (e.g. HEp-2c or MCF-7) were incorporated into the cell culture algorithm also reported preponderance of EV-Cs [8, 9]. It is however important to mention that previous studies in the region that, like the GPLN, used the RD-L20B cell culture based algorithm [19-21] reported a preponderance of EV-Bs. The obvious dissimilarity between the results of these studies might therefore be the cell culture algorithm.

Evidently, the RD-L20B algorithm had selectively filtered out majority of the EV-B containing samples (bearing in mind the EV-B bias of RD cell line [8, 9, 20]), leaving behind more of EV-C containing samples. Hence, it is not unexpected to recover more EV-Cs than EV-Bs in this study. The results of this study therefore add more credence to the EV-B bias hypothesis of RD cell line.

The modifications [16] to the WHO recommended RT-snPCR assays showed that 3.3% (1/30) of the culture negative stool samples from AFP cases analyzed in this study using the WHO recommended RT-snPCR assay had evidence of being mixed (i.e. the case had co-infection with two different enterovirus types). The sample (serial number 1) was shown to contain both EV-B and C (EV-C99 and E25) (Table 1). The inability of the WHO recommended RT-snPCR VP1 assay [11] to detect and resolve mixed isolates has been previously discussed [16]. Hence, as previously suggested [16], the modification of the WHO recommended assay [11] to include independent use of primers 187 and 189 (forward primers) alongside AN88 (reverse primer) for the second round PCR, ensured that the assay retained its sensitivity for enterovirus detection, while providing a simple method to identify and resolve co-infection with different enterovirus types. It is therefore crucial that we do not allow our desire for a panenterovirus detection assay to overshadow our ability to adequately catalogue all enterovirus types present in cases of co-infection.

One of the important isolates detected in this study is a poliovirus serotype 2 (PV-2). It was further shown (Figure 1) that the amino acid at the 143^rd^ position of the VP1 protein was an Isoleucine (Ile_143_), implying that the virus was a vaccine (Sabin) strain [22]. Why this virus was missed by the WHO recommended RD-L20B cell culture based algorithm [5, 6] is currently not clear. Possibilities include break in reverse cold-chain enroute the laboratory and the consequent arrival of the samples with nonviable virus. It is also possible that this PV-2 might represent one of those described by Arita et al., [23] that cannot use the poliovirus receptor (PVR), which is a prerequisite for replication in RD and L20B cell lines. Hence, the inability of the cell lines to detect it. Whatever be the case, the finding of PV-2 in a sample declared negative for such highlights the value of the WHO recommended RT-snPCR assay for detection of enteroviruses even in cases where the enterovirus present might not be detectable by the WHO recommended RD-L20B cell culture based algorithm [5, 6]. This therefore demonstrates the need to use the assay to screen samples declared negative by the RD-L20B cell culture based algorithm before samples are completely ruled as truly negative for poliovirus. The significance of this to the poliovirus eradication effort cannot be over emphasized.

Among all the enterovirus types detected in this study, EV-A119 and EV-B111 are being described for the first time in Nigeria. The significance of this first find of EV-A119 in a child with a clinical condition, particularly AFP, and its zoonotic potential has been described [24]. As regards EV-B111, as at the 4^th^ of March 2017, only three publicly available sequence data of EV-B111 (KF312882, JX538160 and JX538130) existed in GenBank and they were all from South-East Asia. This might therefore represent its first description in Africa. The previously described EV-B111s were recovered in year 2000 in China (KF312882) [25] and 2008 in Bangladesh (JX538160 and JX538130) [26], from humans (KF312882 and JX538160) [25, 26] and nonhuman primates (NHPs) (JX538130) [26]. Consequently, the zoonotic potential of EV-B111 cannot also be ruled out. The fact that only the year 2000 isolate recovered in China was isolated in culture might suggest that EV-B111does not replicate readily in the cell lines used routinely for cell culture in most enterovirology laboratories. Consequently, it is and will mostly be detected by cell culture independent methods like the WHO recommended RT-snPCR assay [11] used in this and the other study [26] that detected it. On the other hand, as evidenced by its varying serology ten years after its description in different prefectures of the same region in China, it is also likely that EV-B111 is not well established in the human population and thus not circulating efficiently [25]. However, as tempting as this proposition is, the fact that EV-B111 has been found at least four times and on two different continents over the last 16 years is course for concern. Furthermore, if EV-B111 is not well adapted to humans, then where has it been surviving and circulating between detections. More troubling is the fact that besides its finding in NHPs [26] the other strains detected in humans till date were found in the stool of children diagnosed with AFP [25, 26]. Of course, the evidence might not be suggestive of causality and might just be indicative of a sensitive AFP surveillance system. In spite these, the occurrence gives reason for further investigation.

## CONFLICT OF INTERESTS

The authors declare that no conflict of interests exist. In addition, no information that can be used to associate the isolates analyzed in this study to any individual is included in this manuscript.

## ACKNOWLEDGEMENTS

We thank the WHO National Polio Laboratory in Ibadan, Nigeria for providing the anonymous samples analyzed in this study. This study was funded by contributions from the authors.

## AUTHOR CONTRIBUTIONS

1. Study Design (JAA, MOA, TOCF)
2. Sample Collection, laboratory and Data analysis (All authors)
3. Wrote, revised, read and approved the final draft of the Manuscript (All authors)

